# Genome assembly of the polyclad flatworm *Prostheceraeus crozieri*

**DOI:** 10.1101/2022.07.12.499707

**Authors:** Daniel J. Leite, Laura Piovani, Maximilian J. Telford

## Abstract

Polyclad flatworms are widely thought to be one of the least derived of the flatworm classes and, as such, are well placed to investigate evolutionary and developmental features such as spiral cleavage and larval diversification lost in other platyhelminths. *Prostheceraeus crozieri*, formerly *Maritigrella crozieri*, is an emerging model polyclad flatworm that already has some useful transcriptome data but, to date, no sequenced genome. We have used high molecular weight DNA extraction and long read PacBio sequencing to assemble the highly repetitive (67.9%) *P. crozieri* genome (2.07 Gb). We have annotated 43,325 genes, with 89.7% BUSCO completeness. Perhaps reflecting its large genome, introns were considerably larger than other free-living flatworms, but evidence of abundant transposable elements suggests genome expansion has been principally via transposable elements activity. This genome resource will be of great use for future developmental and phylogenomic research.

## Introduction

Platyhelminthes (flatworms) are a phylum of protostomes related to annelids, molluscs, and other Lophotrochozoa; they are a very diverse phylum represented by both free-living (turbellarian) and parasitic species (Egger, et al. 2015; Martin-Duran, et al. 2012). They have received particular attention due in part to their parasitism but also to the remarkable regenerative abilities of many species. Members of most flatworm classes are unusual amongst Lophotrochozoa in that they display divergent embryogenic processes (notably Blastomeren Anarchie) that have captured the interests of evolutionary and developmental biologists (Egger, et al. 2015; Martin-Duran, et al. 2012). The canonical spiral cleavage, typical of many Lophotrochozoan phyla, is only seen in the early diverging flatworm classes – Catenulida, Macrostomida, Lecithoepitheliata and Polycladida. Ciliated larvae, comparable to those of annelids and molluscs are even more restricted, being found only in the polyclads. The polyclad class is thus pivotal to understanding the starting point for the evolution of the divergent developmental modes in other platyhelminth classes and more generally for linking platyhelminth development to the wider context of the Lophotrochozoa (Egger, et al. 2015).

*Prostheceraeus crozieri* (previously *Maritigrella crozieri*) is a species of polyclad flatworms found in the mangroves of Bermuda and the Florida Keys. The adults live on (and eat) colonies of the sea squirt species *Ecteinascidia turbinata* (Lapraz, et al. 2013). *P. crozieri* is becoming a useful laboratory model polyclad and transcriptomes of different developmental stages exist; the species has been used to examine early spiral cleavage and larval development using micro-injection labelling techniques, 3D light sheet microscopy (Girstmair and Telford 2019) and gene expression in its Müller’s larva using anti-body and in-situ hybridisation techniques (Rawlinson, et al. 2019).

While previous work has resulted in an assembled *de novo* transcriptome (Lapraz, et al. 2013), a genome is needed to enable comparisons with existing genomes of other free-living flatworms such as the laboratory models *Schmidtea mediterranea* (Grohme, et al. 2018), *Macrostomum lignano* (Wasik, et al. 2015; Wudarski, et al. 2017) and *Dugesia japonica* (An, et al. 2018) as well as those of the many parasitic species. Flatworm genomes are notoriously repetitive and challenging to assemble, but long read sequencing has been used to improve assembly contiguity (Grohme, et al. 2018; Wudarski, et al. 2017).

We have used high molecular weight DNA extracted from a single individual and sequenced with PacBio technology to assemble a draft genome. The genome assembly and annotation will be a key resource for future studies involving this polyclad flatworm.

## Results and Discussion

### The large genome of *P. crozieri*

High molecular weight DNA was extracted from a single, hermaphrodite *P. crozieri* adult and sequenced using PacBio and Illumina technologies, generating 11,921,195 PacBio reads with an N50 of ∼30 kb and 558,509,539 Illumina 150 bp paired end reads, which FastQC identified high quality reads throughout.

The initial assembly used Flye (Kolmogorov, et al. 2019) to assemble PacBio reads to 2.26 Gb, with 26,131 scaffolds and an N50 of 261,667 bp (table 1). Polishing and purging of possible haplotype associated duplicate scaffolds generally removed smaller scaffolds (fig. 1A), reducing the final genome size to 2.07 Gb, with 17,074 scaffolds (16,926 scaffolds >1000 bp) and increased the N50 to 292,050. The assembled genome has a GC content of 37.64% (table 1).

**Table 1.** Genome assembly metricsx

|  | **Initial** | **Polished** | **Purged dups** |
| --- | --- | --- | --- |
| **Size (Gb)** | 2.26 | 2.27 | 2.07 |
| **No. of scaffolds** | 26,131 | 26,131 | 17,074 |
| **N50** | 261,667 | 260,241 | 292,050 |
| **Largest** | 2,607,816 | 2,612,272 | 2,612,272 |
| **N count** | 39,200 | 8,484 | 12,175 |

**Fig. 1.**
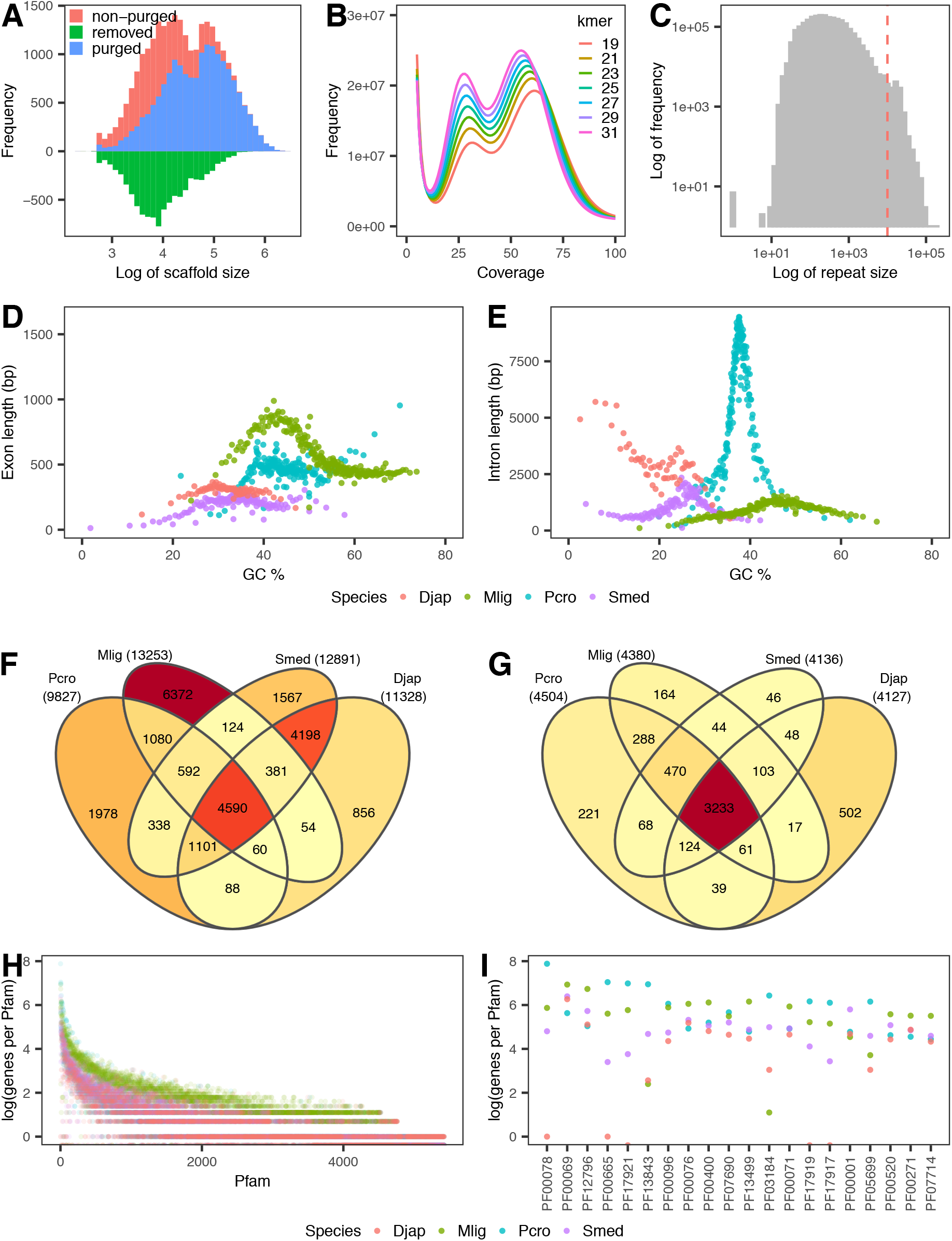
Genome stats, gene annotation characteristics, gene ortholog and Pfam comparison to other free-living flatworms. (**A**) Scaffold size frequency of initial (red) and final assembly (blue) and the scaffold sizes removed (green) during duplicate scaffold removal. (**B**) Kmer frequency coverage reveals two peaks, suggesting diploidy. (**C**) Repeat sizes in the soft-masked genome shows many short and long repeats (>10 kb = red dash line). (**D**) Exon and (**E**) intron sizes and GC% distribution reveal large intron sizes but comparable GC% to other free-living flatworms. Exons/introns were sorted by GC %, split into bins of 1000 genes, and the average length of each bin was measured. (**F**) Orthofinder detected 23,378 orthogroups of which 4,590 (19.6%) were share between all four species. (**G**) Of the total 5,428 Pfams, 3,233 (59.6%) were share between all four species. (**H**) The most abundant Pfam domains ordered by the total of all four species. Mlig in blue shows different distribution relating to possible high gene duplication. (**I**) The top twenty families in (**B**) reveal that *P. crozeri* has a high occurrence of retroviral /transposable element functioning Pfams. Pcro = *P. crozieri* (blue), Smed = *S. mediterranea* (purple), Djap = *D. japonica* (blue) and Mlig = *M. lignano* (green).

This assembled genome is larger than any other free-living flatworm genome known (*S. mediterranea* - 782.1 Mb, *D. japonica* - 1.46 Gb and *M. lignano* - 764 Mb) (An, et al. 2018; Grohme, et al. 2018; Wudarski, et al. 2017). The assembled genome size corresponds closely to a flowcytometry based estimated of 2.5 Gb, indicating an approximately 83% complete assembly (Lapraz, et al. 2013). Kmer-based genome size estimates gave a smaller size of only 1.56 – 1.68 Gb genome size (supplementary table S1), suggesting that Flye performed well despite issues with repeats presumably disrupting kmer based size estimation. Kmer frequencies suggested diploidy, with two peaks occurring (fig. 1B) and predicted heterozygosity levels between 0.810 - 0.936% (supplementary table S1).

The level of duplicate BUSCO genes in the initial assembly was 5.5% and, after polishing and haplotype purging, this was reduced to 2.7% (supplementary table S2). In both assembly versions the percentage of missing BUSCO genes was similar, at ∼13.5% (supplementary table S2), indicating that haplotype specific scaffold removal did not reduce genome completeness.

### Highly repetitive genome

A total of 67.9 % of the genome was identified as repeat and this portion was masked. This level of repeats was a considerable fraction, but this was anticipated given other highly repetitive flatworm genomes (An, et al. 2018; Grohme, et al. 2018; Wasik, et al. 2015; Wudarski, et al. 2017) and the predicted size of this genome. The percent of repeat content was greater than *S. mediterranea* (61.7%), but less than the estimated 80% in *D. japonica*. While retroelements (10.19%) and DNA transposons (23.89%) like PiggyBac and hobo-activator, and SINE (Penelope) and LTR (Pao and Copia), and 1.62% of other repeats (e.g. small RNA, satellites, rolling circles, simple repeats), were identified in the genome, the largest fraction of repeats was unclassified (32.3%).

There were many large repeat regions greater than 10 kb but small repeats were also abundant (fig. 1C). Sequencing and assembly of other free-living flatworms has proved difficult due to the highly repetitive genomes and long repeats, and we also encountered assembly difficulties here, despite using PacBio long reads, likely due to high repeat content and long repeats.

### Many gene annotations have large introns

Braker2 (Bruna, et al. 2021) was used to predict gene models and predicted a total of 43,325 genes, with 46,235 isoforms, which had an average length of 2,048 bp. 23,852 of the 43,325 genes had transcriptional support >1 transcript per million (TPM) in the RNAseq data. InterProScan (Jones, et al. 2014) identified 21,493 of the predicted genes with homology to Pfam domains and, of these, 12,199 were also supported by the existing transcriptome data. This suggests that Braker2 was able to recover gene predictions that had Pfam homology but which lacked RNAseq evidence. The BUSCO completeness of the annotated gene set [C:89.7% [S:87.1%, D:2.6%], F:5.2%, M:5.1%] was more complete than the genome assembly alone (Table 3).

We compared the length and GC content of exons and introns with other free-living flatworms (Zhu, et al. 2009). *P. crozieri* exons had a mean length of 467 bp, which was similar to what is seen in *S. mediterranea* (198 bp), *D. japonica* (297 bp) and *M. lignano* (574 bp) (fig. 1D). However, *P. crozieri* introns were substantially longer than what is seen in the three other flatworms, with *P. crozieri* having an average intron length of 5,263 bp compared to *S. mediterranea* (1,064 bp), *D. japonica* (2,972 bp) and *M. lignano* (975 bp) (fig. 1E). *P. crozieri* average exon GC content was 44.5% (higher than the genome GC of 37.64%), which was greater than *S. mediterranea* and *D. japonica*, but less than *M. lignano* (fig. 1D). The GC of introns (37.4%) was very similar to the background *P. crozieri* genomic GC content (fig. 1E).

### Comparisons of Pfam domain content with other flatworms

Orthofinder (Emms and Kelly 2019) analysis identified 23,378 orthogroups of which 4,590 orthogroups were shared between *P. crozieri, S. mediterranea, D. japonica* and *M. lignano* (fig. 1F). Many orthogroups were shared between the closely related *S. mediterranea* and *D. japonica* (4,198) or found only in *M. lignano* (6,372) (fig. 1F).

Across all four species, a total of 5,428 Pfams were detected, with 3,233 being shared in all four species (fig. 1G). We also asked how many genes were associated with each Pfam domain in the other available free-living flatworm genomes. The number of genes per Pfam domain was similar in *P. crozieri, S. mediterranea* and *D. japonica* but the macrostomid *M. lignano* had more instances of genes linked to each Pfam, supporting previous evidence of high levels of duplication in *M. lignano* (fig. 1H) (Wasik, et al. 2015; Wudarski, et al. 2017). It is possible that the large number of specific orthology groups in *M. lignano* is associated with the divergence of these duplicated genes (Holland, et al. 2017; Natsidis, et al. 2021).

Many of the most frequently occurring Pfam domains in *P. crozieri* (rvt_1 [pf00078], rve [pf00665], piggybac [pf13843] and integrase [pf17921]), were also more abundant than the other flatworms (fig. 1I) and are associated with retroviral or transposable element genes. Taken together with the high proportion of repetitive elements it could suggest that *P. crozieri* has a large number of active transposable elements. It is unclear whether the large intron sizes (when compared to other flatworms), are functionally related to the higher transposable element activity.

### Homeobox gene repertoire

We annotated 89 homeobox containing genes in *P. crozieri* (29 ANTP, 19 PRD, 11 LIM, 7 TALE, 6 SINE, 4 POU, 3 CUT, 3 ZF, 1 CERS, 1 HNF, 2 PROS and 3 unassigned) (supplementary fig. S1, supplementary table S3), which covers the 11 major classes (Holland, et al. 2007), which is similar to other free-living flatworms (Abril, et al. 2010; Currie, et al. 2016; Olson 2008). We found five Hox genes *Hox1, Hox6-8* and three *Hox9-13*/*Post2*. ParaHox genes (*Cdx, Gsx* and *Xlox*/*Pdx*) have been lost (or not identified) in *S. mediterranea* (Currie, et al. 2016); we identified *Cdx* and *Gsx* but not *Xlox*/*Pdx* in *P. crozieri* (supplementary table S3). The Hox genes were not found in a single cluster although two *Hox9-13* genes were linked on a single scaffold, *Cdx* and *Hhex* were present on another scaffold and tandem duplicates of *Otx* on a third (supplementary table S3). Low discovery of syntenic homeobox genes may be a result of a large, repeat-rich genome that is fragmented. While the *P. crozieri* genome is considerably larger than other flatworms sequenced to date, the lack of duplicated homeobox and BUSCO genes implies there have not been large-scale or whole genome duplications in the lineage leading to *P. crozieri*.

### Genes associated with pluripotency and regeneration

Like other flatworms, *P. crozieri* possesses high regenerative capabilities (Lapraz, et al. 2013). Flatworms have lost most mammalian stem cell and pluripotency genes (*Oct4*/*Pou5f1, Nanog, Klf4, c-Myc*, and *Sox2*) however. Of these mammalian factors, only *Sox2* homologs remain in *S. mediterranea* and *M. lignano* (Grohme, et al. 2018; Wasik, et al. 2015). Similarly, in *P. crozieri, Sox2* was present in one copy, and none of the other factors were identified indicating that, despite its regenerative capabilities, *P. crozieri* lacks the same genes as other flatworms.

## Conclusion

We have assembled and annotated the first polyclad flatworm genome of *P. crozieri* attaining a 2.07 Gb assembly with 43,325 genes. The high repeat content of 67.9 % was not unexpected based on other flatworm genomes. Despite the problems that these large repeat contents can cause in genome assembly, high BUSCO scores and the homeobox repertoire suggests the assembly and annotation are of reasonable completeness and quality that will be useful for future studies. Our work helps elevate *P. crozieri* as an increasingly important model that will contribute to our understanding of flatworm and animal evolution.

## Materials and Methods

### Animal collection, DNA extraction and sequencing

*P. crozieri* adults were collected between Largo and Marathon Key from the Florida Keys, USA (September/October 2019), transported in sea water to UCL, UK and transitioned to artificial sea water (ASW) and maintained in ASW for four weeks. DNA from one live adult was extracted following a standard soft tissue protocol from BioNano Prep Animal tissue DNA Isolation. Extracted DNA was stored at 4°C for three days before DNA concentration was estimated using NanoDrop and TapeStation technology. Approximately 10 μg of DNA used for library preparation and sequencing with two SMRT SQII PacBio cells and shearing, library preparation and 150 bp paired-end Illumina sequencing at University of California, Berkeley, CA, USA.

### Kmer genome size estimation

Genome size was estimated with kmer abundance in short read data with Jellyfish v2.3 (Marcais and Kingsford 2011) using kmer lengths of 21, 23, 25, 27, 29, 31 bp, with option count -C. Histo generated files using Jellyfish histo were used with GenomeScope (read_length=150, kmer_max=10,000) to estimate the genome size and heterozygosity (Vurture, et al. 2017) and visualised with R v3.5.3.

### Genome assembly

We use the repeat concatenated de Bruijn graph assembler Flye v2.7 (Kolmogorov, et al. 2019) and the PacBio reads for an initial assembly with the genome size parameter set to 2.5 Gb (-g 2.5g), 75x coverage for repeat graph construction (--asm-coverage 75) and a minimum overlap of 8,000 bp (-m 8000) to avoid an overly fragmented assembly. This was followed by one round of polishing with long reads using Flye (Kolmogorov, et al. 2019).

Further polishing with NextPolish v1.1.0 (Hu, et al. 2020) short reads trimmed with Trimmomatic v0.39 (LEADING:3 TRAILING:3 SLIDINGWINDOW:4:15 MINLEN:36) (Bolger, et al. 2014). long reads to polish using the -task=best strategy. The parameters for minimap2 v2.17-r941 (Li 2018) for max depth of short reads was set to 35x coverage and for long reads -x map-pb, with a minimum read length of 5 kb, maximum read length 300 kb and max depth at 60x.

Purge_dups v1.2.3 (Guan, et al. 2020) further collapsed haplotype scaffolds (including parameter -e). We searched for BUSCO genes at each step of assembly and the final gene predictions. Busco v3.0.2 (Simao, et al. 2015) was used with metazoan_odb9 with default evalue and “-long” for optimisation of the Augustus parameters in genome searches.

### Repeat modelling and masking

*De novo* repeats were identified with RepeatModeler v2.0.1 (Flynn, et al. 2020), with RepeatScout v1.0.6 (Price, et al. 2005), TandemRepatsFinder v4.06 (Benson 1999) and RECON v1.08 (Bao and Eddy 2002), Genometools v1.6 ltrharvest (Ellinghaus, et al. 2008; Gremme, et al. 2013), LTR_retriever v2.8 (Ou and Jiang 2018), with the RMBlast v2.10.0 search engine and the -LTRstruct identification options. This *de novo* repeat library and the Dfam3.2 (Hubley, et al. 2016) library were used with RepeatMasker v4.0.7 to produce a soft masked genome assembly of *P. crozieri*.

### Gene prediction and annotation

For gene annotation we used RNA-seq evidence with the Braker v2.1.2 (Bruna, et al. 2021) pipeline with Augustus v3.2.3 (Stanke, et al. 2006) and GeneMark-ET v4.46 (Bruna, et al. 2020). First, the RNAseq data from PRJNA275060 included paired end (SRR1801815) and single end (SRR1801812) data were trimmed with Trimmomatic v0.39 (Bolger, et al. 2014) (LEADING:3 TRAILING:3 SLIDINGWINDOW:4:15 MINLEN:36). The soft-masked genome was indexed with Star v2.7.3a (Dobin, et al. 2013) and reads were mapped using the multi-sample 2-pass method to improve accuracy of splice junction information. BAM files were sorted by coordinates with Samtools v1.9 (Li, et al. 2009) as RNAseq evidence for Braker v2.1.2 (Bruna, et al. 2021) to predict gene models including their UTRs (-UTRs=on), using 10 rounds of optimisation (-r 10) and CRF modelling (-crf). Interproscan v 5.47-82.0 (Jones, et al. 2014) was used to annotate protein predictions with all available databases. These Interproscan results, along with Interproscan searches for *S. mediterranea, M. lignano* and *D. japonica*, were used to assess Pfams in free-living flatworm and presence of pluripontency genes (*Nanog, Klf4, c-Myc*, and *Sox2*) in *P. crozeri*.

### Homeobox gene annotation

The homeodomain PF00046 Pfam RP55 alignment was used with hmmsearch v3.3.1 (Eddy 2011) to query the *P. crozieri* protein annotations and domain hits were extracted using esl-sfetch v0.47. Hits (length > 50 amino acids) were aligned with all *Caenorhabditis elegans, Branchiostoma floridae* and *Tribolium castaneum* homeodomains from HomeoDB (Zhong, et al. 2008; Zhong and Holland 2011) (http://homeodb.zoo.ox.ac.uk/) using MAFFT v7.475 with 1000 iterations (Katoh and Standley 2013). Iqtree v2.0.3 (Minh, et al. 2020) built maximum likelihood trees, using 1000 ultrafast bootstraps with automatic model prediction (LG+G4). The consensus tree was visualised in Figtree.

## Data availability

All genomic sequence data has been deposited under the BioProject PRJEB44148. The genome assembly has been uploaded to ENA (GCA_907163375) and annotations and a brief description of the assembly and annotation pipeline have been made accessible at https://github.com/djleite/PROCRO_genome.

## Acknowledgements

This work was supported by a Leverhulme Trust Research Project [grant number RPG-2018- 302 to MJT and DJL] and by the European Union’s Horizon 2020 research and innovation programme under the Marie Skłodowska-Curie grant agreement no. 766053 [EvoCELL: grant to MJT, fellowship to LP]. We also thank Johannes Girstmair and Florida Keys Marine Lab for their help in animal collection, and Martin Tran for their help with high molecular weight DNA extractions.

## Supplementary Tables

**Supplementary table S1.**
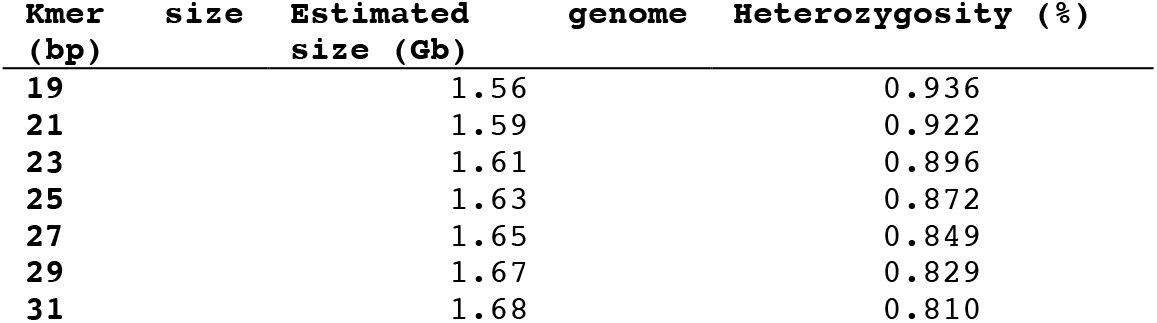
GenomeScope genome size and heterozygosity estimations at different Kmer sizes

**Supplementary table S2.** Percentage BUSCO scores for assembly stages and gene annotation

|  | Initial | Polished | Purged dups | Soft-masked | Gene annotation |
| --- | --- | --- | --- | --- | --- |
| Complete | 79.2 | 79.8 | 79.0 | 79.9 | 89.7 |
| Single | 73.7 | 76.3 | 75.8 | 77.2 | 87.1 |
| Duplicate | 5.5 | 3.5 | 3.2 | 2.7 | 2.6 |
| Fragmented | 7.3 | 6.3 | 7.7 | 6.4 | 5.2 |
| Missing | 13.5 | 13.9 | 13.3 | 13.7 | 5.1 |

**Supplementary table S3.**
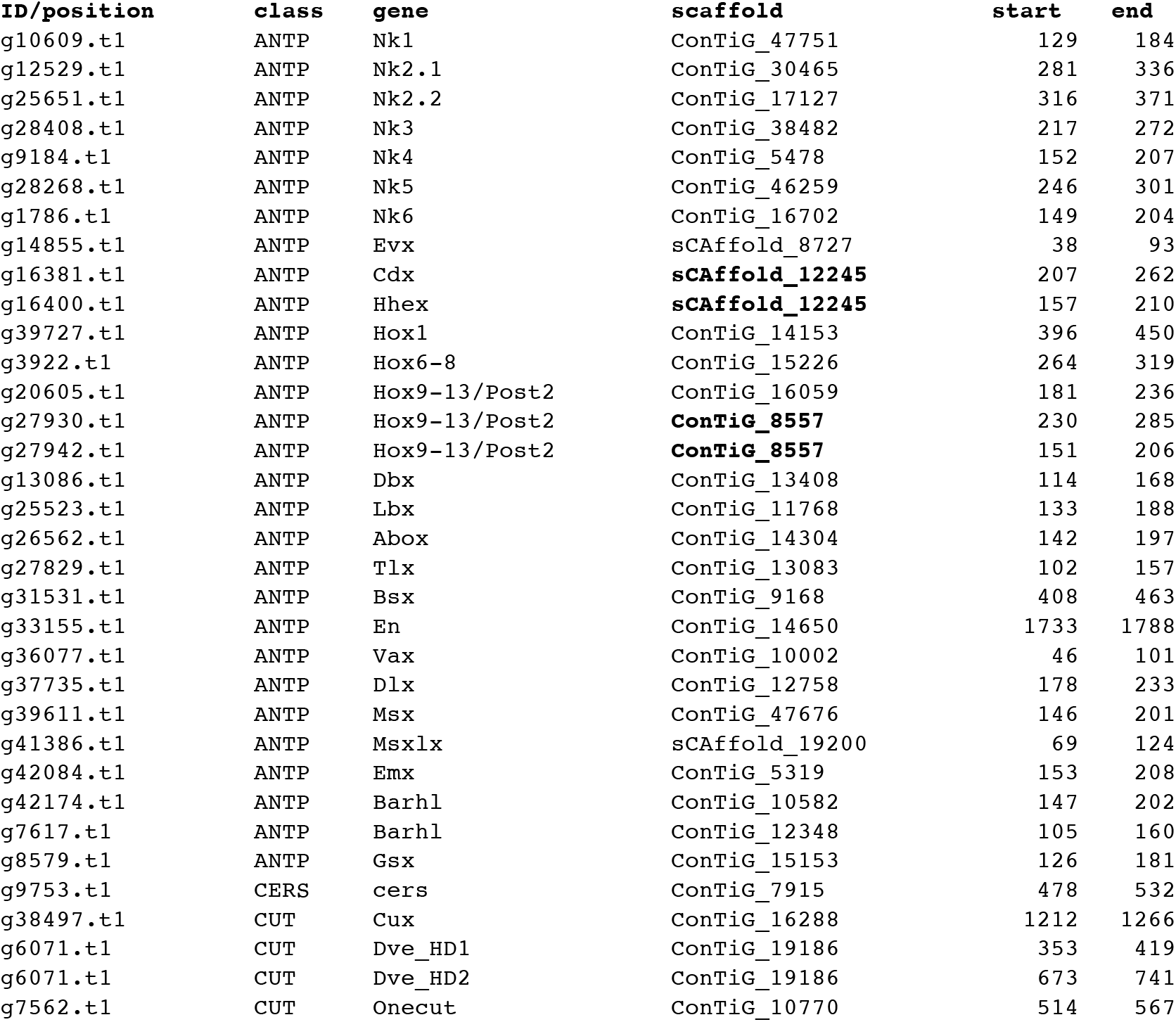
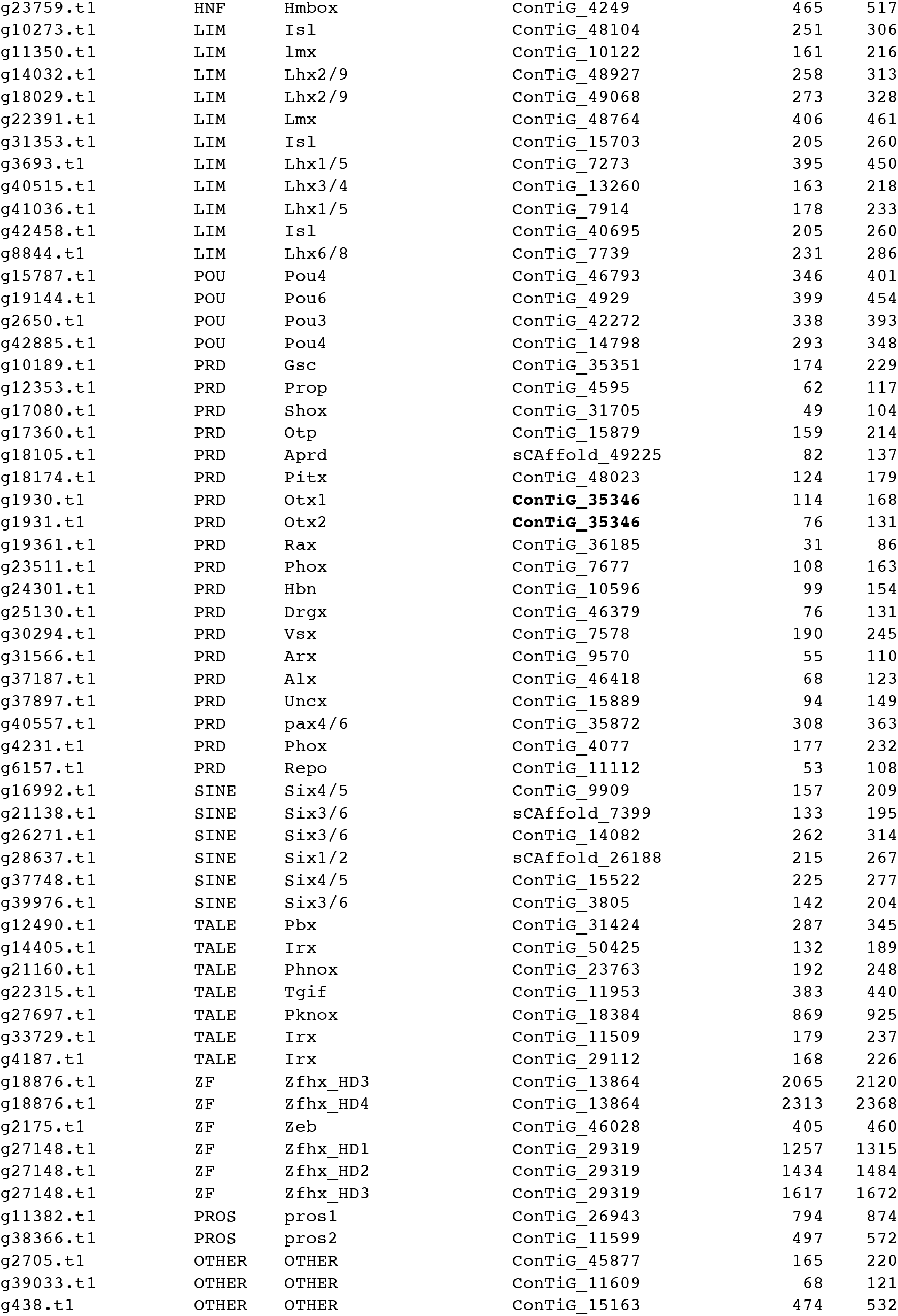
Homeobox annotations in *P. crozieri*.

## Supplementary figures

**Supplementary Fig. S1.**
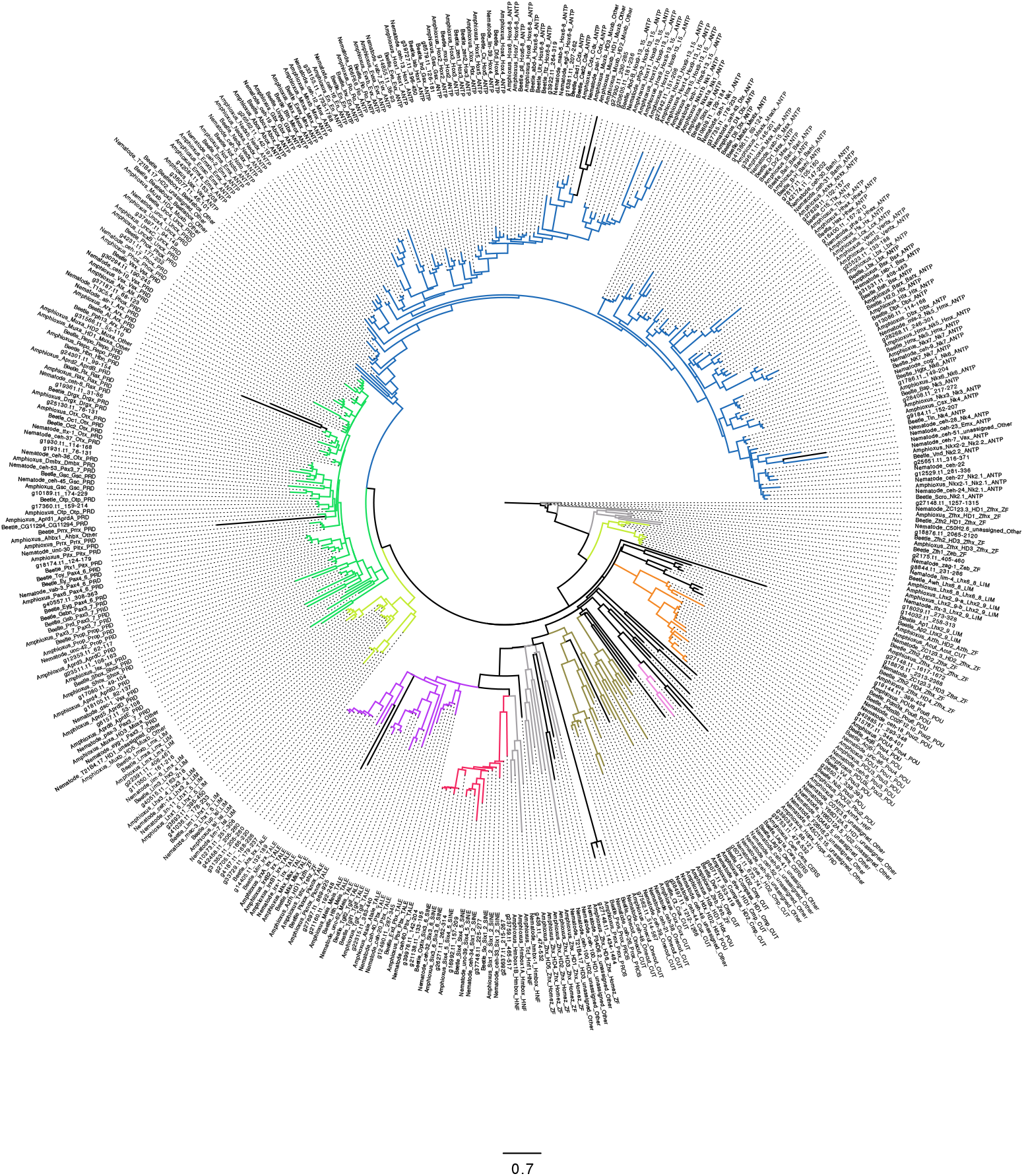
Maximum likelihood tree reconstruction of *P. crozeri* homeodomains. Eleven major classes of homeobox genes were identified in *P. crozeri*, PROS class not shown in the tree. Annotations were made based on their relationship to *C. elegans*, Amphioxus and *T. castaneum* homeodomains from HomeoDB. LG+G4 model with 1000 ultrafast bootstraps.

